# Protein characterization of the soybean malic enzyme family to select metabolic targets for seed oil improvement

**DOI:** 10.1101/2025.03.18.644018

**Authors:** Mariana Beatriz Badia, Joaquín Costa, Juan Ignacio Zucchetti, Tatiana Pavlovic, Paula Calace, Mariana Saigo, Mariel Claudia Gerrard Wheeler

**Author notes:** Corresponding author: Mariel Claudia Gerrard Wheeler, Suipacha 570, S2000LRJ, Rosario, Argentina, Email address https://www.cefobi-conicet.gov.ar/grupo-gerrard-wheeler/. Instituto de Agrobiotecnología Rosario (INDEAR) (Mariana Beatriz Badia). Instituto de Biología Molecular y Celular de Rosario (IBR-CONICET). These authors have contributed equally to this work and should be considered co-first authors (Mariana Beatriz Badia, Joaquín Costa).

## Abstract

Seeds are of great economic interest due to their organic composition. The metabolic pathways leading to the deposition of seed storage compounds, such as oil and proteins, are complex in terms of the number of enzymes involved, the regulatory context, and the variability among species. Previous studies indicated that mitochondrial NAD-dependent and plastidial NADP-dependent malic enzymes (MEs; EC 1.1.1.39 and 1.1.1.40, respectively) participate in providing reducing power and citrate or acetyl-CoA, respectively, to fatty acid biosynthesis in soybean (*Glycine max* L. Merr.) embryos. In this work, we analyzed the biochemical properties of enzymes belonging to the soybean ME family; which were obtained by recombinant expression and subsequent affinity purification. Our results show that NADP-ME1.1 has high affinity for the substrates malate and NADP and is specifically activated by glutamine, the main nitrogenous compound imported from maternal tissues to support the filling process. In addition, NAD-MEs exhibit differential kinetic parameters and regulation by key metabolites, making the isoforms better suited to fulfill non-redundant functions under certain physiological conditions. Activation by intermediates of glycolysis, the citric acid cycle, and lipogenesis indicated coordination between these pathways and malate metabolism. In turn, NAD-ME activity in soybean mitochondria would be regulated by the alternative association between linage 1 and 2 subunits. These findings provide new insights into intermediary metabolism in oilseeds and nominate specific soybean ME isoforms as biotechnological tools to be used in breeding programs.

## 1. Introduction

Seeds are essential for the reproduction of flowering plants, as they contain, protect, and nourish the developing embryo (Baud and Lepiniec, 2010). The storage compounds accumulated during seed maturation are important raw materials for both human and animal food and the energy industry (Tokel and Erkenciogu, 2021). Typically, soybean seeds contain, in addition to proteins and carbohydrates, about 20% triacylglycerols (Kambhampati et al., 2021). Although its oil content is lower than that of other oil plant species such as castor (60%), sunflower (50%), or rapeseed (40%), soybean is one of the most versatile and important agronomic crops worldwide. The fatty acid composition of soybean oil has been altered by metabolic engineering to improve its utility and nutritional quality (Haun et al., 2014; Al Amin et al., 2019). However, attempts to modify the proportion of reserves have not been entirely successful, possibly due to the complexity and poor understanding of the pathways of seed metabolism (Arias et al., 2022; Aznar-Moreno et al., 2022; Morley et al., 2023).

Fatty acid synthesis for seed triacylglycerols occurs in plastids and utilizes carbon skeletons (acetyl-CoA), energy (ATP), and reduction equivalents (NAD(P)H) generated from maternal nutrients (Sagun et al., 2023). The glycolytic conversion of sucrose to phosphoenolpyruvate (PEP) followed by the action of PEP carboxylase and malate dehydrogenase or pyruvate kinase results in the synthesis of cytosolic malate or pyruvate, respectively (Hajduch et al., 2011). Both metabolites are then imported into the plastid and transformed into acetyl-CoA by the action of the pyruvate dehydrogenase complex, following the conversion of malate to pyruvate by a NADP-dependent malic enzyme (NADP-ME; EC 1.1.1.40). In this regard, a single transcript encoding a plastidial NADP-ME (NADP-ME1.1; Glyma13g43130) has been detected in soybean embryos, and its accumulation corresponds to the developmental stages during which oil deposition is most active (Gerrard Wheeler et al., 2016). Furthermore, NADP-ME1.1 would produce reducing power directly where fatty acids biosynthesis occurs.

In addition, a high contribution of amino acids to fatty acids synthesis has been described in soybean (Allen et al., 2009; Allen and Young, 2013), probably related to the plant’s ability to form nitrogen-fixing nodules. Amino acids are mainly degraded in mitochondria, providing significant amounts of tricarboxylic acid cycle intermediates. This contribution becomes important especially from the R6.5 stage onwards, where carbohydrates start to support the generation of protective compounds that help during final seed desiccation (Gerrard Wheeler et al., 2016; Aznar-Moreno et al., 2022). In this context, the NAD-dependent malic enzyme (NAD-ME; EC 1.1.1.39) enables the relocation of mitochondrial carbon to the plastid for seed filling, a process that is mediated by citrate transport (Allen and Young, 2013; Pavlovic et al., 2023). As soybean seed matures, a large increase in NAD-ME activity was reported, which correlates with the accumulation of NAD-ME1, NAD-ME2.1, NAD-ME2.3, and NAD-ME2.4 transcripts (Glyma03g24630, Glyma09g39870, Glyma03g01680, and Glyma07g08110, respectively; Gerrard Wheeler et al., 2016). At this point, these isoforms would provide pyruvate, and thus acetyl-CoA, to promote mitochondrial citrate synthesis.

Therefore, the above results nominate the ME gene family as a key target for soybean improvement. However, a comprehensive characterization of the protein products is necessary to develop accurate biotechnological strategies (Bar-Even et al., 2013). In this work, the kinetic, structural, and regulatory properties of soybean MEs were determined and comparatively analyzed in order to understand the rate and specificity of carbon flux during seed filling and select the best targets to generate genotypes with higher oil content.

## 2. Results

### 2.1 Kinetic parameters indicate different catalytic performances of soybean NAD- and NADP-ME isoforms

The soybean ME isoforms to be characterized were selected on the basis of their predicted subcellular localization and expression patterns (Gerrard Wheeler et al., 2016; Table S1). NAD-ME1 (Glyma03g24630), NAD-ME2.1 (Glyma09g39870), NAD-ME2.3 (Glyma03g01680), NAD-ME2.4 (Glyma07g08110), and NADP-ME1.1 (Glyma13g43130) increase their transcript levels as embryo maturation progresses, participating in precursor input for seed reserve biosynthesis. NADP-ME1.1 is a plastidial protein belonging to the NADP-ME family, whereas the NAD-ME counterparts are putative mitochondrial proteins. Among the latter, NAD-ME1 corresponds to lineage 1 and NAD-ME2.1, NAD-ME2.3, and NAD-ME2.4 correspond to lineage 2, each evolutionarily defined by At2g13560 (AtNAD-ME1) and At4g00570 (AtNAD-ME2) of *Arabidopsis thaliana*, respectively (Gerrard Wheeler et al., 2016; Tronconi et al., 2018).

High sequence conservation is observed among members comprising NAD-MEs lineages 1 and 2 and among NADP-utilizing enzymes (Table S1). Thus, soybean NAD-ME1 shares 83% and 87% identity with its Arabidopsis and castor orthologs, whereas it showed lower identity (65%) with soybean type 2 enzymes (84-97% identity between them; Table S1). In this context, AlphaFold structure models showed that NAD-ME2.4 and NAD-ME2.3 are highly similar, while root mean squared deviation and template modeling score of NAD-ME2.4 and NAD-ME1 alignment indicated more differences between these structures (Table S2; Figure S1; Videos 1 and 2). Although three phylogenic subgroups are distinguished in soybean NADP-MEs (Gerrard Wheeler et al., 2016; Arias et al., 2018), they are very similar (77-98% identity), compared to NAD-dependent isoforms (37-42% identity; Table S1).

*E. coli* strains transformed with the vectors containing the cDNA encoding soybean ME isoforms were able to overexpress mature proteins. Each protein was found in the soluble fraction of the bacterial extract and subsequently purified to acceptable quantity and purity. The monomeric molecular masses estimated by SDS-PAGE agreed with *in silico* predictions: 66 kDa for NAD-ME2.3 and NAD-ME2.4, 67 kDa for NAD-ME1 and NAD-ME2.1, and 70 kDa for NADP-ME1.1 (Figure S2).

First, the malate decarboxylation reaction was recorded at different pHs (between 4.5 and 8.9) to establish the subsequent assay conditions. The pH conditions with the maximum activities for each ME isoform were identified (Table 1). Thus, we selected pH 6.5 and 7.5 for the NAD-ME and NADP-ME assays, respectively. The NAD-ME isoenzymes only showed detectable activity in the presence of MnCl_2_, whereas NADP-ME1.1 was active with both MnCl_2_ and MgCl_2_, showing higher activity values with the former metal cation (Table 1).

**Table 1.** Kinetic parameters of soybean NAD- and NADP-ME isoforms. Values are the mean of three different experiments with the corresponding standard deviation. ND: not detected. -: not determined. NAD-ME isoforms showed no detectable reverse activity. ^1^S_0.5 pyruvate_ (mM), ^2^nH _piruvate_, ^3^k_cat_/S_0.5 pyruvate_ (s^-1^ mM^-1^).

| REACTION | FORWARD |  |  |  |  | REVERSE |
| --- | --- | --- | --- | --- | --- | --- |
| ENZYME | NAD-ME1 | NAD-ME2.1 | NAD-ME2.3 | NAD-ME2.4 | NADP-ME1.1 |  |
| Optimum pH | 6.5-8.0 | 5.5-6.5 | 6.5 | 5.5-6.5 | 7.0-8.0 | 6.5 |
| Activity with MnCl<sub>2</sub>/MgCl<sub>2</sub> (U ml<sup>-1</sup>) | 73.8 ± 18.0/ND | 9.4 ± 4.1/ND | 10.0 ± 1.5/ND | 0.5 ± 0.1/ND | 428.4 ± 18.3/269.8 ± 78.9 | 107.0 ± 25.7/83.7 ± 20.1 |
| k<sub>cat</sub> (s<sup>-1</sup>) | 265.3 ± 18.0 | 122.4 ± 6.0 | 102.8 ± 6.9 | 65.5 ± 7.3 | 211.5 ± 8.9 | 202.4 ± 6.9 |
| S<sub>0.5</sub> malate (mM) | 0.3 ± 0.1 | 3.3 ± 0.3 | 3.2 ± 0.2 | 5.1 ± 1.2 | 0.6 ± 0.1 | 5.1 ± 0.5<sup>1</sup> |
| nH malate | 1.2 ± 0.4 | 1.4 ± 0.2 | 1.7 ± 0.2 | 1.2 ± 0.2 | 0.8 ± 0.1 | 1.1 ± 0.1<sup>2</sup> |
| k<sub>cat</sub>/S<sub>0.5</sub> malate (s<sup>-1</sup> mM<sup>-1</sup>) | 884.3 | 37.1 | 32.1 | 12.8 | 352.5 | 39.7<sup>3</sup> |
| S<sub>0.5</sub> NAD (mM) | 0.4 ± 0.1 | 1.3 ± 0.3 | 13.8 ± 3.9 | 6.8 ± 1.5 | - | - |
| S<sub>0.5</sub> NADP (μM) | - | - | - | - | 5.6 ± 0.7 | - |
| nH NAD(P) | 1.2 ± 0.3 | 1.1 ± 0.2 | 0.9 ± 0.1 | 1.0 ± 0.1 | 1.3 ± 0.2 | - |

The kinetic parameter determinations indicated that NAD-ME1 has higher k_cat_ and substrate affinities than the others isoforms using NAD as a cofactor (Table 1). Thus, the S_0.5_ values for malate of NAD-ME2.1, NAD-ME2.3, and NAD-ME2.4 are 11- to 17-fold higher than those of NAD-ME1. Like NAD-ME1, NADP-ME1.1 also showed a high k_cat_ and low S_0.5_ for malate (Table 1). Furthermore, we observed a large difference with respect to the affinity for the dinucleotide cofactor, with a S_0.5 NADP_ value for NADP-ME1.1 equal to 5.6 µM, compared to S_0.5 NAD_ values ranging from 0.4 to 13.8 mM for the NAD-ME isoforms. We determined hyperbolic behavior for the soybean enzymes (nH close to unity), except for NAD-ME2.3 with malate, for which a sigmoidal response was observed (nH = 1.7; Table 1).

### 2.2 Some metabolites differentially affect enzymatic activity depending on the isoform

Next, we determined the effect of several metabolites on the activity of soybean ME isoforms. The results showed that each enzyme exhibited different behavior towards a particular compound; essentially, the overall response of the lineage 2 NAD-dependent isoforms was different from that of NAD-ME1 or NADP-ME1.1 (Figure 1). Proteins of lineage 2 showed positive regulation by intermediates produced in irreversible steps of the glycolytic pathway, such as fructose-1,6-bis-P and PEP (effect observed in NAD-ME2.1, NAD-ME2.3, and NAD-ME2.4) and glucose-6-P (in the case of NAD-ME2.4). In addition, these isoforms showed strong activation by CoA and acetyl-CoA (Figure 1). Lineage 2 NAD-ME isoforms also specifically increased their activity in the presence of the organic acid oxaloacetate. In contrast, NAD-ME1 showed only minor activation with fructose-1,6-bis-P and inhibition with oxaloacetate (Figure 1). In the case of NADP-ME1.1, we observed specific activation by the amino acid glutamine (Figure 1). Likewise, citrate activated NAD-ME2.1, NAD-ME2.4, and NADP-ME1.1 without affecting the other isoforms (Figure 1).

**Figure 1.**
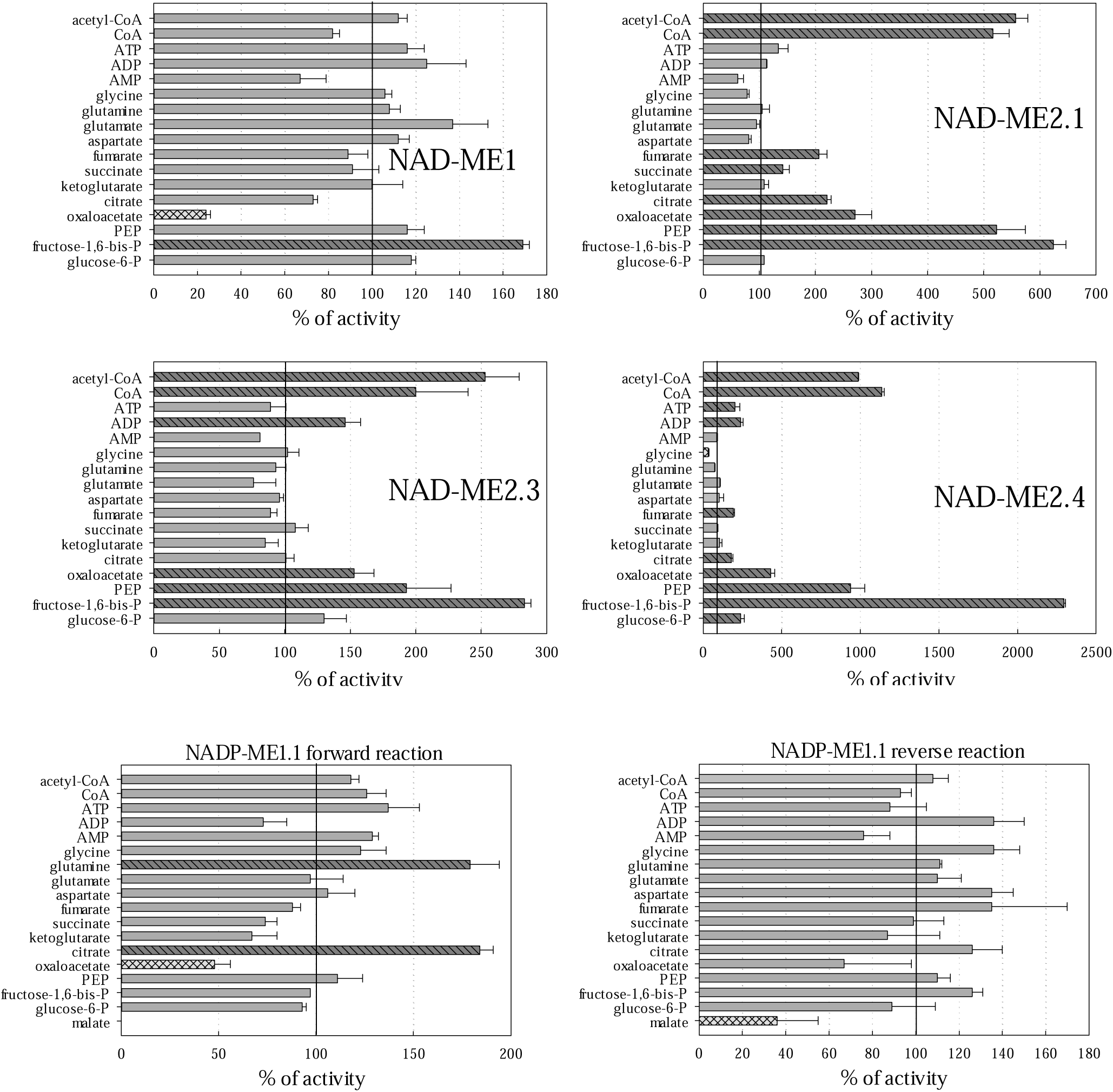
Regulatory properties of soybean NAD- and NADP-ME isoforms. Data are presented as ratio (in percentages) of the activity in the presence of each metabolite compared to a control in the absence of the metabolite. Determinations were performed in triplicate (means and standard deviations are showed). Effectors that produced a significant change (Student’s t-test, *p* < 0.05) are shown in light grey and double dashed line (inhibition) or dark grey and dashed line (activation). Assays were performed at pH optimum using sub-saturating levels of substrates. With the exception of CoA and acetyl-CoA (both assayed at 20 μM), metabolites were used at 2 mM. NADP-ME1.1 also catalyzes the reverse reaction so it was assayed for both directions.

We also evaluated the performance of recombinant enzymes in the presence of total polar metabolite extracts from soybean embryos at progressive stages of development; i.e., from early-, mid-, and late-filling embryos (R5.5 to R7) to fully mature dry embryos (R8; Fehr et al., 1971). NADP-ME1.1 exhibited a peak of activation at the onset of filling (Figure S3A) and the profile matches total NADP-ME activity levels in developing embryos (Gerrard Wheeler et al., 2016). In the case of NAD-dependent activity, the pattern (Figure S3B) does not match activity measurements, where a continuous increase is observed throughout R5.5 to R8 (Gerrard Wheeler et al., 2016). This result is likely due to the contribution of multiple isoforms to the total activity, with different expression levels and kinetic parameters (Table 1; (Gerrard Wheeler et al., 2016). Still, the assay with metabolite extracts from embryos revealed that NAD-ME2.1 is activated by the mixture present in embryos at the end of filling (140-190% of activity; R6.5 and R7 in Figure S3B). In turn, NAD-ME2.3 was activated throughout the range of stages (190-300%); whereas no substantial modifications of NAD1-ME activity were observed (70-125%). Finally, NAD-ME2.4 showed a trend towards inactivation at the end of embryo development (25-55 % activity; Figure S3B).

### 2.3 Only NADP-ME1.1 is able to catalyze the reductive carboxylation of pyruvate

We also evaluated the ability of soybean ME isoforms to catalyze the reverse direction of the reaction, i.e. reductive carboxylation of pyruvate. NADP-ME1.1 was able to catalyze the reverse reaction in the presence of NADPH, with both MgCl_2_ (83.7 U ml^-1^) and MnCl_2_ (107.0 U ml^-1^; Table 1). In contrast, NAD-dependent isoforms did not catalyze pyruvate carboxylation under any of the conditions tested.

Next, we determined the kinetic parameters of the NADP-ME1.1 reverse activity. An optimum pH of 6.5 was determined, a more acidic value than for the forward reaction, i.e. oxidative decarboxylation of malate (Table 1). The k_cat_ for both directions were similar. However, the affinity of the enzyme for pyruvate was lower than for malate (higher S_0.5_ in Table 1). Thus, the catalytic efficiency (k_cat_/S_0.5_) of the reverse reaction was 9-fold lower than that of the forward reaction. We did not observe any positive regulation for the NADP-ME1.1 reverse activity with any metabolite; only malate generated an inhibition by reaction product (Figure 1).

### 2.4 The kinetic and structural properties of NAD-ME2.3 and 2.4 isoforms are affected by the addition of NAD-ME1

Based on the ability of AtNAD-ME2 to form a hetero-dimer with AtNAD-ME1 in Arabidopsis cells [19, 20], we mixed the NAD-ME2.3 or NAD-ME2.4 isoforms with NAD-ME1 and evaluated the kinetic parameters and regulatory properties of each protein mixture. The k_cat_ values were 1.3 or 1.5-fold higher than that of NAD-ME1 alone (Table 2). Although both mixtures showed similar affinities toward malate and NAD substrates compared to NAD-ME1 (Tables 1 and 2), the response of metabolic effectors also differed in terms of individual NAD-ME activities (Figure 2). Interestingly, NAD-ME2.3 in the presence of NAD-ME1 gained regulation by metabolites such as fumarate, succinate, ketoglutarate, citrate, and glucose-6-P; whereas it lost activation by CoA/acetyl-CoA and oxaloacetate, characteristic of lineage 2 NAD-ME isoforms (Figure 1). Moreover, the NAD-ME1 + NAD-ME2.4 mixture is only activated by CoA, glutamate, and ketoglutarate (Figure 2).

**Figure 2.**
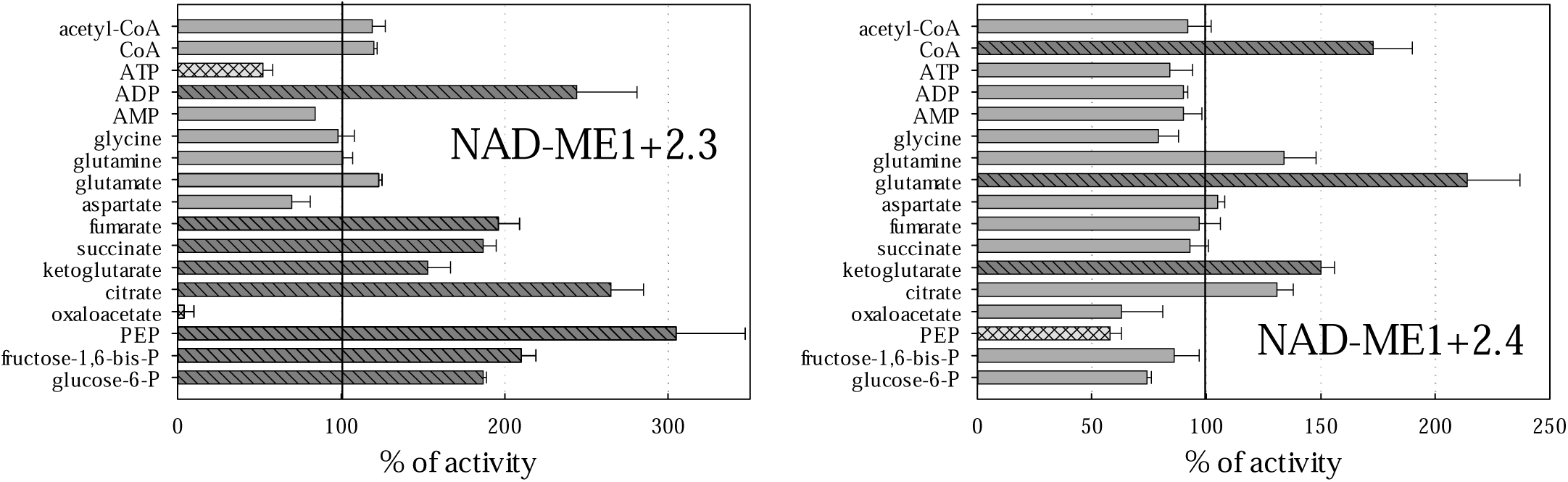
Regulatory properties of NAD-ME mixtures. Data are presented as ratio (in percentages) of activity in the presence of each metabolite compared to a control in the absence of the metabolite. Determinations were performed in triplicate (means and standard deviations are showed). Effectors that produced a significant change (Student’s t-test, *p* < 0.05) are shown in light grey and double dashed line (inhibition) or dark grey and dashed line (activation). Assays were performed at pH 6.5 using sub-saturating levels of malate and NAD. With the exception of CoA and acetyl-CoA (both assayed at 20 μM), metabolites were used at 2 mM.

**Table 2.** Kinetic parameters of NAD-ME equimolar mixtures. Values are the mean of three different experiments with the corresponding standard deviation. The asterisk indicates a significant difference (Student’s t-test, *p* < 0.05) with respect to the value obtained for the enzymes separately (Table 1).

|  | NAD-ME1 + NAD-ME2.3 | NAD-ME1 + NAD-ME2.4 |
| --- | --- | --- |
| $k_{\text{cat}}$ ($\text{s}^{-1}$) | $338.3 \pm 15.8^*$ | $393.6 \pm 63.8^*$ |
| $S_{0.5 \text{ malate}}$ (mM) | $0.2 \pm 0.1$ | $0.6 \pm 0.2$ |
| $nH_{\text{malate}}$ | $1.1 \pm 0.1$ | $0.9 \pm 0.2$ |
| $k_{\text{cat}}/S_{0.5 \text{ malate}}$ ($\text{s}^{-1} \text{ mM}^{-1}$) | 1,691.5 | 656.0 |
| $S_{0.5 \text{ NAD}}$ (mM) | $0.2 \pm 0.1$ | $0.2 \pm 0.1$ |
| $nH_{\text{NAD}}$ | $0.8 \pm 0.3$ | $1.2 \pm 0.3$ |
| $k_{\text{cat}}/S_{0.5 \text{ NAD}}$ ($\text{s}^{-1} \text{ mM}^{-1}$) | 1,691.5 | 1,968.0 |

In addition, principal component analysis of kinetic data suggested differences between the NAD-ME isoforms and that mixing them generates new entities (Figure 3A). Dimension 1 opposes NAD-ME1 and NAD-ME2.4 to the mixture NAD-ME1+ NAD-ME2.4 (Figure 3A; strongly positive and negative coordinates on the x-axis, respectively). NAD-ME2.4 clusters with the more closely related NAD-ME2.1 and NAD-ME2.3 (Figure 3A, graph of individuals), forming a group characterized by activation by glycolytic intermediates and fatty acid precursors (Figure 3A, graph of variables). In turn, dimension 2 opposes NAD-ME1+ NAD-ME2.3 to the individual components and also of NAD-ME1 + NAD-ME2.4 (Figure 3A, strongly positive and negative coordinates on the y-axis, respectively). Both clusters differ mainly in the high or low values of variables related to the regulation of enzymatic activity by metabolites (Figure 3A).

**Figure 3.**
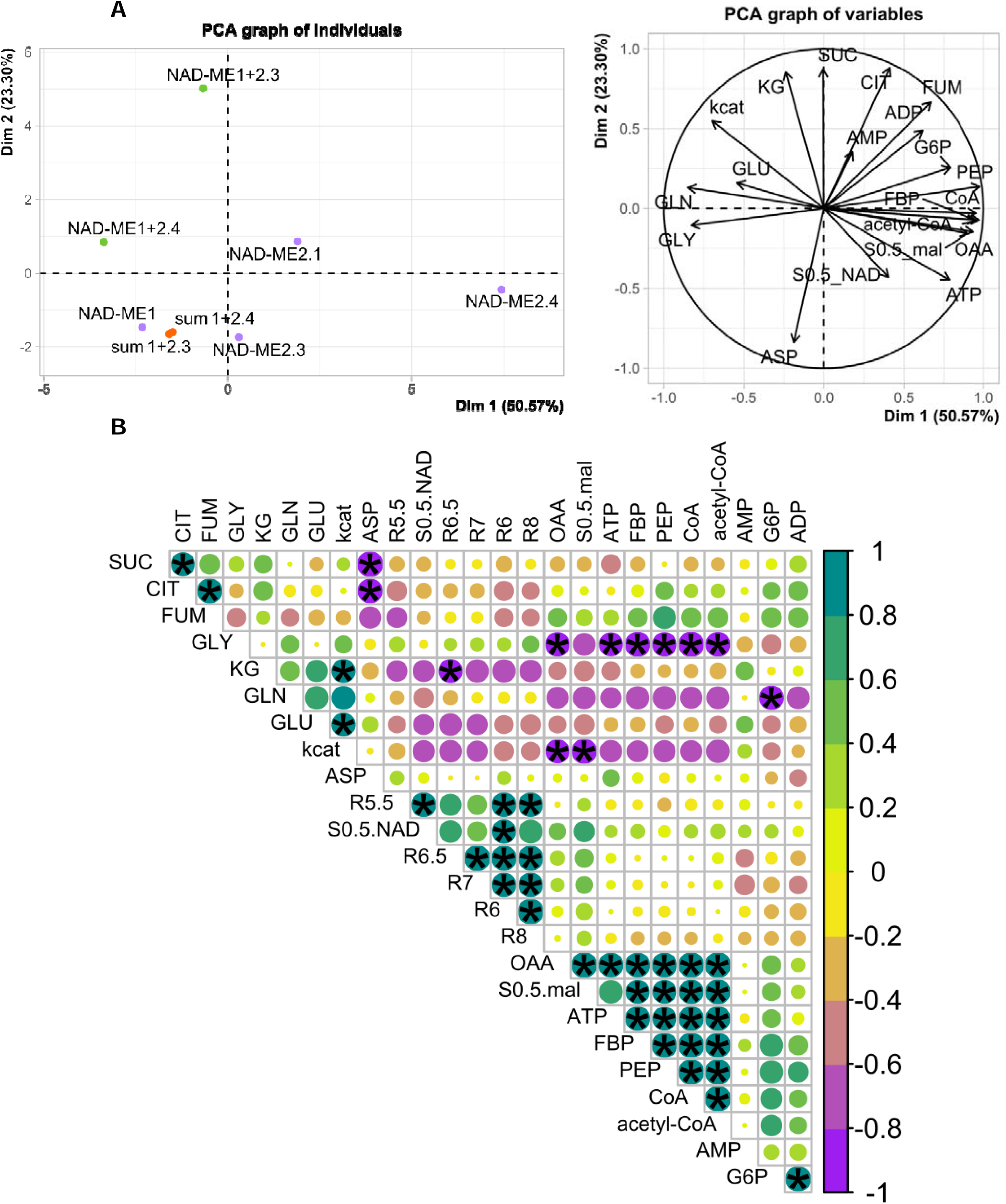
Kinetic data analysis of soybean NAD-ME isoforms. **A**) Principal component analysis (PCA) including 8 individuals and 20 variables (from Figures 1 and 2 and Tables 1 and 2). Dimensions (Dim) 1 and 2 account for a high percentage of the variability in the data (73.87%). The sum 1 + 2.3 (or 2.4) is the theoretical value assuming no interaction for the isoform mixture between NAD-ME1 plus NAD-ME2.3 or NAD-ME2.4. **B)** Correlation plot including also the results with mixtures of embryo metabolites (Figures 1 and 2 and Tables 1 and 2 plus Figure S3). The color of the circles indicates the correlation between variables, with 1 and −1 being the maximum positive and negative correlations, respectively. The size of the circles represents the degree of significance of the correlation, with an asterisk for that with *p* < 0.05. GLY: glycine, GLN: glutamine, GLU: glutamate, ASP: aspartate, FUM: fumarate, SUC: succinate, KG: ketoglutarate, CIT: citrate, OAA: oxaloacetate, FBP: fructose-1,6-bis-P, G6P: glucose-6-P, MAL: malate.

### 2.5 Relationship between kinetic parameters and metabolic regulation of NAD-ME isoforms

To identify statistical relationship between kinetic data for NAD-ME isoforms, we performed a correlation analysis. The correlation plot detected groups of metabolites with similar effects and interactions between kinetic parameters (Figure 3B). We observed a high positive correlation between the activation of NAD-MEs by oxaloacetate, ATP, fructose-1,6-bis-P, PEP, CoA, and acetyl-CoA, and in turn with a high value of S_0.5malate_ (Figure 3B). Something similar occurred in relation to the affinity for NAD, since the enzymes most activated by metabolites extracted from developing embryos were those with the highest S_0.5NAD_ (Figure 3B). On the other hand, a negative correlation was observed between k_cat_ and S_0.5_ plus oxaloacetate regulation, probably related to the mechanism of catalysis (Figure 3B). Finally, the regulation of NAD-ME isoforms by citric acid cycle intermediates appears to be coordinated and functionally related (Figure 3B).

## 3. Discussion

Soybean plants produce and store triacylglycerols in the seeds, useful as food/feed and energy renewable sources. Therefore, improving lipid content and composition has generated immense research efforts in order to select targets for metabolic engineering. ME can be used (Morley et al., 2023) into a strategy ("push strategy"; Sagun et al., 2023) to increase the amount of precursors (pyruvate and NAD(P)H in this case) flowing into fatty acid biosynthetic pathways. Carbon allocation is usually regulated by different mechanisms, including transcriptional control of enzyme-encoding genes, subcellular compartmentalization, and inhibition of divergent pathways to minimize secondary products (Arias et al., 2022; Aznar-Moreno et al., 2022). In addition, kinetic parameters and allosteric regulation of key enzymes play a critical role in this fine control (Hartman et al., 2023; Kroll et al., 2023). The most sought-after targets are those that have a high catalytic efficiency and that are not negatively regulated by seed metabolites. Understanding these properties can lead to the design of more successful transgenic events; and in this work, we performed biochemical studies to assess the catalytic function and metabolic control of soybean ME isoforms induced during storage compound deposition in embryos. There is no report on the analysis of mitochondrial and plastidial members of the soybean ME family; although a cytosolic NADP-ME has been associated with tolerance to aluminum stress, contributing to the metabolism and efflux of malate and citrate in soybean roots (Zhou et al., 2018). In other previous work, Morley et al. (2023) constructed and analyzed homozygous transgenic soybean plants expressing plastidial AtNADP-ME4 (Gerrard Wheeler et al., 2005) or mitochondrial AtNAD-ME2 (Tronconi et al., 2008) from Arabidopsis. Alterations in central metabolism that result in a subtle but significant increase in oil content (2-9%) were observed in AtNADP-ME4 events, whereas the AtNAD-ME2 transgene produced no differences from wild-type seed oil (Morley et al., 2023). Both proteins have been extensively characterized in our laboratory; in particular, AtNADP-ME4 showed a lower k_cat_ (151.3 s^-1^; Gerrard Wheeler et al., 2005) than soybean NADP-ME1.1 (Table 1) and a tendency to be inhibited by key metabolic intermediates (oxaloacetate, fumarate, ATP, and glucose-6-P; Gerrard Wheeler et al., 2008). Furthermore, AtNAD-ME2 was assayed alone and not in combination with AtNAD-ME1 (Morley et al., 2023), with the hetero-dimer being the entity with the highest activation tendency (Tronconi et al., 2008; 2010). The kinetic parameters and regulatory capacity of soybean ME isoforms are discussed below and some of them are nominated as new improved targets to obtain genotypes with higher seed quality.

Soybean NAD-dependent isoforms do not carboxylate pyruvate; whereas NADP-ME1.1 is capable of catalyzing this reverse reaction, albeit with lower affinity for the substrate than the forward reaction (estimated from S_0.5_ values; Table 1). Furthermore, we observed negligible modulation of the reverse activity of NADP-ME1.1 by key metabolites (Figure 1). These results support that NAD(P)-ME isoforms would function in vivo primarily in the direction of oxidative decarboxylation of malate.

In contrast, NAD(P)-ME forward activity is modulated by several compounds involved in different metabolic pathways (Figure 1). Citrate was postulated as an important metabolite involved in carbon allocation to maternal organic acid in plastidial biosynthesis of seed storage compound precursors (Allen et al., 2009; Allen and Young, 2013; Pavlovic et al., 2023). In this regard, we observed a positive modulation of NADP-ME1.1 activity by citrate (Figure 1). In addition, NADP-ME1.1 showed a specific activation by glutamine, the main nitrogenous compound imported into the soybean embryo from the photosynthetic tissues of the mother plant (Egli and Bruening, 2001). Thus, this plastidial isoform would be more active when the embryo synthesizes reserves, i.e. when maternal nutrients are available and there is an enhanced transport of carbon and nitrogen to the site where it is located. Glycolytic intermediaries activated lineage 2 NAD-ME isoforms (Figure 1), suggesting an increase in pyruvate and NADH production in coordination with a block in glycolysis (Plaxton and Podestá, 2006). Likewise, the concomitant activation of NAD-ME2.1, NAD-ME2.3 and NAD-ME2.4 by CoA and acetyl-CoA (Figure 1) suggests their role in the supply of precursors for fatty acid biosynthesis, as well as plastidial NADP-ME from castor bean endosperm (Shearer et al., 2004). The non-hyperbolic response to malate in NAD-ME2.3 (Table 1) revealed that the substrate itself may also be a regulator of enzymatic activity in a cooperative manner.

Soybean NAD-ME1 exhibited lower regulation (Figure 1) but better performance, compared to the plant isoforms characterized to date (Tronconi et al., 2008; Calace et al., 2021; Hüdig et al., 2022). This enzymatic fitness is based on both the highest k_cat_ and lowest S_0.5_ values, resulting a catalytic efficiency (Table 1) more than 8-fold higher than the “average enzyme” (100 s^-1^ mM^-1^, obtained from data on several thousand enzymes of different organisms collected in the literature; (Bar-Even et al., 2011). Another isoform that exceeds this threshold is plastidial NADP-ME1.1 (Table 1). Despite the similarity of the structures of NAD-ME2.3 and NAD-ME2.4 (Figures S1 and S2; Tables S1 and S2), their biochemical properties and allosteric regulation are somewhat different (Table 1; Figures 1 and S3). Thus, subtle structural changes affecting ligand binding or signal transduction induce fine-tunning of NAD-ME activity and highlight that it is not possible to define, based on sequence data alone, which enzyme will have the desired characteristics.

Soybean NAD-ME isoforms showed enzymatic activity, each separately: NAD-ME1 presents a better kinetic fitness (Table 1), whereas lineage 2 isoforms have a better ability to be regulated by pure or seed-extracted metabolites (Figures 1 and S3). We also observed changes in the properties of mixtures containing NAD-ME1 plus NAD-ME2.3 or NAD-ME2.4 (Tables 1 and 2; Figures 1 and 2), probably due to the formation of new entities. The kinetic parameters and regulation by metabolites do not correspond to the mere sum of enzymes (Figure 3A), indicating that NAD-ME activity in soybean mitochondria would be regulated by alternative association between subunits. In this regard, the increase in the k_cat_ for NAD-ME1 + NAD-ME2.3 and NAD-ME1 + NAD-ME 2.4 (Table 2) together with the high capacity of NAD-ME1 + NAD-ME2.3 to be activated by a large number of metabolites (Figure 2) is noteworthy.

Taken together, the findings reported here provide new insights into intermediate metabolism in soybean embryos and nominate the highly active and metabolite-activated proteins, in particular NADP-ME1.1 and/or the oligomers of NAD-ME1 plus NAD-ME2.3, as promising new biotechnological targets for plant breeding or oil production in other simpler systems.

## 4. Materials and Methods

### 4.1 Plant material

Soybean (*Glycine max* L. Merr.) Pioneer 94m80 was grown in greenhouse, in soil pots supplemented weekly with 0.5X Hoagland solution, under natural sunlight plus metal halide illumination to provide a 16-h light regime and temperatures between 20°C (night) and 30°C (day). Soybean plants were sorted according to reproductive stages (R5.5-R8; Fehr et al., 1971), and seeds were harvested and dissected to separate embryos, which were stored at −80°C.

### 4.2 Cloning of soybean NAD(P)-ME cDNAs

Total RNA was isolated from 100 mg of each embryo sample using TRIzol reagent, purified on PureLink spin columns (Thermo Fisher Scientific), and treated with RQ1 DNase (Promega). RNA was converted into first-strand cDNA using SuperScriptII Reverse Transcriptase (Invitrogen) and oligo (dT)_15_ primer (Biodymamics). Full-length cDNAs were amplified by PCR from R7 RNA for NAD-ME2.1, NAD-ME2.3, and NAD-ME2.4; and from R8 RNA for NAD-ME1 and NADP-ME1.1. Phusion DNA polymerase (Thermo Fisher Scientific) and isoform-specific primers were used. These primers were designed to introduce unique NheI or NdeI and XhoI sites at the 5′- and 3′-ends, respectively. For expressing mature proteins, the 5′-primers included the first codon downstream of the predicted transit peptide cleavage site (TargetP 1.1 server; http://www.cbs.dtu.dk/services/TargetP/) and the 3′-primers contained the stop codon. The following primer combinations were used: NAD-ME1<u>NheI</u> (5′-GCTAGCCCCTCCATCGTCCACA-3′) and NAD-ME1<u>XhoI</u> (5′-CTCGAGTCATTCTTTCTTGTAAACTAA-3′), NAD-ME2.1<u>NdeI</u> (5′-CATATGTTATCGACGGCGATTC-3′) and NAD-ME2.1<u>XhoI</u> (5′-CTCGAGCTATTTTTCATGAACAAGAGG-3′), NAD-ME2.3<u>NheI</u> (5′-GCTAGCTGCATTGTTCACAAGCGCGGTGCCGACATACTCCACGAT-3′) and NAD-ME2.3<u>XhoI</u> (5′-CTCGAGTTATTTTTCATGAACGAGAGG-3′), NAD-ME2.4<u>NheI</u> (5′-GCTAGCTGCATTGTTCACAAGCGCGGTGCCGACATACTCCATGAC-3′) and NAD-ME2.4<u>XhoI</u> (5′-CTCGAGTCATTTTTCATGAACGAGAGG-3′), and NADP-ME1.1<u>NheI</u>(5′-GCTAGCAGTATGACCCCAAGCA-3′) and NADP-ME1.1<u>XhoI</u>(5′-CTCGAGTCATCGGTAGCTTCTGTAGGC-3′). The amplified products were introduced into pBluescript II SK (+) (Stratagene) by blunt-end restriction and ligation procedure. NAD(P)-ME sequences were confirmed by full-length sequencing using generic primers (T3 and T7 promoter) at Macrogen service.

### 4.3 Generation of vectors for protein expression

Soybean NAD(P)-ME cDNAs contained in pBluescript II SK (+) were digested with NheI (or NdeI) and XhoI (Promega) for subcloning into the expression vector pET28 (Novagen). Thus, only 19-21 amino acids from the vector are added to the protein sequence. The structure of the resulting plasmids was verified by restriction enzyme analysis. The molecular mass of each protein was predicted by Expasy server (http://www.web.expasy.org/protparam/).

### 4.4 Heterologous expression and affinity purification of recombinant enzymes

Vectors pET28 containing soybean NAD(P)-ME cDNAs were used to express the recombinant proteins fused in-frame to a His-tag using *Escherichia coli* BL21(DE3) as host. The transformed cells were grown in 200 ml LB medium in the presence of 50 µg ml^-1^ kanamycin until each culture reached A_600_ = 0.8. Lactose 0.5% (w v^-1^) was added as an inducer and the cells were further cultured for 16 h at 15°C. Cells were then collected by centrifugation for 10 min at 10,000 x g and resuspended in buffer containing 20 mM Tris-HCl (pH 7.5), 0.25 M NaCl, 0.01% (v v^-1^) Triton X-100, and 1 mM phenylmethylsulfonyl fluoride.

After sonication to achieve cell lysis and subsequent centrifugation, each soluble fraction was used for protein purification on a His-NTA column (Novagen) containing nickel, according to the protocol described by Saigo et al., (2013). Purified enzymes were desalted and concentrated on Centricon YM-50 devices (Millipore) using 50 mM Tris-HCl (pH 7.5), 10 mM MnCl_2_ (or MgCl_2_), and 50% (v v^-1^) glycerol. Integrity and purity were analyzed by SDS-PAGE (Laemmli, 1970).

### 4.5 Enzymatic activity and protein concentration assays

NAD-ME forward activity was assayed using standard reaction mixtures containing 10 mM MnCl_2_, 10 mM malate, and 4 mM NAD in a final volume of 0.5 ml (Tronconi et al., 2018). The standard medium for NADP-ME forward activity assays contained 10 mM MgCl_2_, 10 mM malate, and 0.5 mM NADP (Gerrard Wheeler et al., 2005). Enzyme activities were recorded at different pHs as follows: 50 mM sodium acetate-acetic acid (pH 4.5-5.5), 50 mM MES-NaOH (pH 6.0-6.5), 50 mM MOPS-KOH (pH 7.0), and 50 mM Tris-HCl (pH 7.5-8.9). According to the results, pH 6.5 and 7.5 were selected for the NAD-ME and NADP-ME assays, respectively. Reductive carboxylation of pyruvate (reverse reaction) was assayed in 50 mM MES-NaOH pH 6.5, 10 mM MgCl_2_, 30 mM pyruvate, 50 mM NaHCO_3_, and 0.2 mM NADPH (Gerrard Wheeler et al., 2008). Each reaction was initiated with the addition of each enzyme or an equimolar mixture of them. One unit (U) is defined as the amount of enzyme that catalyzes the formation (or consumption) of 1 μmol of NAD(P)H per min under the specified conditions (ε_340nm_ = 6.22 mM^−1^ cm^−1^). All assays were performed at 30°C in a Jasco V-730 spectrophotometer. The detection limit was less than 0.02 nmol of NAD(P)H formed (or consumed) per min.

Kinetic parameters were determined by varying the concentration of one substrate while maintaining the other substrate(s) at saturation level, at the optimum pH of the reaction. Substrate solutions were prepared with MnCl_2_ or MgCl_2_ to avoid sequestration of the essential metal cofactor. The following K_d_ values were used: K_d_ (Mn-malate) = 20 mM, K_d_ (Mn-NAD) = 12.9, K_d_ (Mg-malate) = 28.2 mM, and K_d_ (Mg-NADP) = 19.1 mM (Grover et al., 1981). Data were analyzed based on free substrate concentrations in the assay media. Substrate-dependent rates were fitted to the Hill equation:

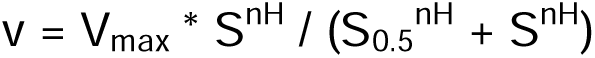

where v represents the initial rate at varying substrate concentration (S), V_max_ is the maximum reaction rate, nH is the Hill index, and S_0.5_ is the substrate concentration for which half of V_max_ is obtained. The catalytic constant (k_cat_) corresponds to the μmol of substrate converted to product per s and μmol of active site under optimal conditions. Protein concentration was determined by Bio-Rad protein assay (Bradford, 1976) using bovine serum albumin as standard.

To identify possible inhibitors or activators, NAD(P)-ME forward activity was measured in the absence or presence of 0.5 or 2.0 mM of different metabolites (glucose-6-P, fructose-1,6-bis-P, PEP, oxaloacetate, citrate, ketoglutarate, succinate, fumarate, aspartate, glutamate, glutamine, glycine, AMP, ADP, and ATP). For CoA and acetyl-CoA, a final concentration of 20 µM was used. For reverse activity, malate was also evaluated as possible effector. The enzymes were also assayed in the presence of total polar metabolite extracts from developing soybean embryos (Gerrard Wheeler et al., 2016). In this case, the amount in each reaction medium was equivalent to the metabolites present in 2 mg of embryo tissue. Substrate levels were always maintained at the S_0.5_ values determined for each isoenzyme.

### 4.6 Structural determinations

Structure models were retrieved from the AlphaFold Protein Structure Database (Jumper et al., 1981; https://alphafold.ebi.ac.uk/) and compared using the Pairwise Structural Aligment Tool (https://www.rcsb.org/alignment).

### 4.7 Statistical analysis

Student’s t-test was used for comparisons between two parameters. Sample size and significance are indicated in the legends. Correlation plot was constructed using the Hmisc and Corrplot packages in R and clustered using the hclust model with *p*< 0.05. Principal components analysis was performed with the Factoshiny package in R.

## Acknowledgements

MS and MCGW belong to the Researcher Career of National Council for Scientific and Technical Research; MBB, TP and PC are fellows of the same institution. JIZ is a fellow of the National Agency for the Promotion of Research, Technological Development and Innovation (Agencia I+D+i). This work was supported by Agencia I+D+i (PICT 2019-02618), Santa Fe Agency of Science, Technology and Innovation (IO-2018-00141), and National University of Rosario (80020180300060UR and 80020220700146UR). The funding sources were not involved in the research or in the preparation and presentation of the article. The authors would like to thank Gina Dotta and Mariana Giró for their technical assistance in carrying out this work.

## Abbreviations

k_cat_: catalytic constant
ME: malic enzyme
NAD-ME: NAD-dependent malic enzyme
NADP-ME: NAD-dependent malic enzyme
nH: Hill index
PCA: principal component analysis
PEP: phosphoenolpyruvate
S: substrate concentration
S_0.5_: substrate concentration for half of V_max_
U: unit of enzymatic activity
v: initial rate
V_max_: maximum reaction rate

## Author contributions

MCGW designed the concept and planned the experiments in this study. MBB and JC performed the cloning and protein expression assays. MBB, JC, PC, and MCGW purified the recombinant enzymes. MBB, JC, JIZ, TP, and MCGW performed the enzymatic activity assays. MS was involved in the statistical analysis and structural determinations. MBB, JC, MS, and MCGW analyzed and interpreted the results. MCGW, in collaboration with the other authors, drafted and edited the manuscript.

