## Supplementary material for "Protein characterization of the soybean malic enzyme family to select metabolic targets for seed oil improvement"

| Soybean enzyme | Gene ID | Subcellular Localization | *Arabidopsis thaliana* | | | | | |
| --- | --- | --- | --- | --- | --- | --- | --- | --- |
|  |  |  | AtNAD-ME1 | AtNAD-ME2 | AtNADP-ME1 | AtNADP-ME2 | AtNADP-ME3 | AtNADP-ME4 |
| GmNAD-ME1^a^ | Glyma03g24630 | mitochondria | 83 | 64 | 42 | 39 | 39 | 38 |
| GmNAD-ME2.1^a^ | Glyma09g39870 | mitochondria | 62 | 78 | 39 | 37 | 38 | 37 |
| GmNAD-ME2.2 | Glyma18g46340 | mitochondria | 62 | 78 | 39 | 37 | 38 | 37 |
| GmNAD-ME2.3^a^ | Glyma03g01680 | mitochondria | 62 | 79 | 39 | 38 | 38 | 39 |
| GmNAD-ME2.4^a^ | Glyma07g08110 | mitochondria | 63 | 81 | 40 | 39 | 40 | 39 |
| GmNADP-ME1.1^a^ | Glyma13g43130 | plastid | 38 | 38 | 78 | 76 | 76 | 77 |
| GmNADP-ME1.2 | Glyma15g02230 | plastid | 38 | 38 | 78 | 77 | 76 | 77 |
| GmNADP-ME1.3 | Glyma08g21530 | plastid | 40 | 39 | 76 | 75 | 75 | 80 |
| GmNADP-ME2.1 | Glyma04g09110 | cytosol | 42 | 38 | 80 | 81 | 80 | 78 |
| GmNADP-ME2.2 | Glyma06g09220 | cytosol | 41 | 38 | 79 | 80 | 80 | 78 |
| GmNADP-ME3.1 | Glyma05g35800 | cytosol | 39 | 37 | 81 | 79 | 78 | 73 |
| GmNADP-ME3.2 | Glyma01g01180 | cytosol | 40 | 38 | 82 | 78 | 77 | 78 |
| GmNADP-ME3.3 | Glyma16g08460 | cytosol | 39 | 37 | 82 | 78 | 78 | 76 |

| Soybean enzyme | Gene ID | Subcellular Localization | *Ricinus communis* | | | | |
| --- | --- | --- | --- | --- | --- | --- | --- |
|  |  |  | RcNAD-ME1 | RcNAD-ME2 | RcNADP-ME1 | RcNADP-ME2 | RcNADP-ME3 |
| GmNAD-ME1^a^ | Glyma03g24630 | mitochondria | 87 | 67 | 38 | 40 | 40 |
| GmNAD-ME2.1^a^ | Glyma09g39870 | mitochondria | 64 | 84 | 38 | 39 | 38 |
| GmNAD-ME2.2 | Glyma18g46340 | mitochondria | 64 | 85 | 38 | 39 | 38 |
| GmNAD-ME2.3^a^ | Glyma03g01680 | mitochondria | 63 | 85 | 38 | 39 | 38 |
| GmNAD-ME2.4^a^ | Glyma07g08110 | mitochondria | 64 | 86 | 37 | 41 | 40 |
| GmNADP-ME1.1^a^ | Glyma13g43130 | plastid | 37 | 39 | 81 | 81 | 82 |
| GmNADP-ME1.2 | Glyma15g02230 | plastid | 37 | 39 | 80 | 81 | 82 |
| GmNADP-ME1.3 | Glyma08g21530 | plastid | 39 | 40 | 84 | 80 | 78 |
| GmNADP-ME2.1 | Glyma04g09110 | cytosol | 40 | 39 | 84 | 86 | 84 |
| GmNADP-ME2.2 | Glyma06g09220 | cytosol | 40 | 38 | 83 | 86 | 83 |
| GmNADP-ME3.1 | Glyma05g35800 | cytosol | 39 | 39 | 78 | 83 | 88 |
| GmNADP-ME3.2 | Glyma01g01180 | cytosol | 40 | 39 | 84 | 82 | 87 |
| GmNADP-ME3.3 | Glyma16g08460 | cytosol | 39 | 38 | 81 | 82 | 88 |

| Soybean enzyme | *Glycine max* | | | | |
| --- | --- | --- | --- | --- | --- |
|  | GmNAD-ME1 | GmNAD-ME2.1 | GmNAD-ME2.2 | GmNAD-ME2.3 | GmNAD-ME2.4 |
| GmNAD-ME1^a^ | - | 65 | 65 | 65 | 65 |
| GmNAD-ME2.1^a^ | 65 | - | 95 | 84 | 86 |
| GmNAD-ME2.2 | 65 | 95 | - | 86 | 87 |
| GmNAD-ME2.3^a^ | 65 | 84 | 86 | - | 97 |
| GmNAD-ME2.4^a^ | 65 | 86 | 87 | 97 | - |
| GmNADP-ME1.1^a^ | 37 | 37 | 37 | 38 | 38 |
| GmNADP-ME1.2 | 37 | 37 | 37 | 38 | 38 |
| GmNADP-ME1.3 | 39 | 38 | 38 | 38 | 40 |
| GmNADP-ME2.1 | 41 | 38 | 38 | 39 | 40 |
| GmNADP-ME2.2 | 40 | 38 | 37 | 38 | 39 |
| GmNADP-ME3.1 | 38 | 38 | 38 | 39 | 38 |
| GmNADP-ME3.2 | 41 | 37 | 37 | 38 | 39 |
| GmNADP-ME3.3 | 39 | 36 | 37 | 38 | 38 |

| Soybean enzyme | *Glycine max* | | | | | | | |
| --- | --- | --- | --- | --- | --- | --- | --- | --- |
|  | GmNADP-ME1.1 | GmNADP-ME1.2 | GmNADP-ME1.3 | GmNADP-ME2.1 | GmNADP-ME2.2 | GmNADP-ME3.1 | GmNADP-ME3.2 | GmNADP-ME3.3 |
| GmNAD-ME1^a^ | 37 | 37 | 39 | 41 | 40 | 38 | 41 | 39 |
| GmNAD-ME2.1^a^ | 37 | 37 | 38 | 38 | 38 | 38 | 37 | 36 |
| GmNAD-ME2.2 | 37 | 37 | 38 | 38 | 37 | 38 | 37 | 37 |
| GmNAD-ME2.3^a^ | 38 | 38 | 38 | 39 | 38 | 39 | 38 | 38 |
| GmNAD-ME2.4^a^ | 38 | 38 | 40 | 40 | 39 | 38 | 39 | 38 |
| GmNADP-ME1.1^a^ | - | 97 | 87 | 81 | 81 | 77 | 80 | 77 |
| GmNADP-ME1.2 | 97 | - | 87 | 82 | 81 | 77 | 80 | 77 |
| GmNADP-ME1.3 | 87 | 87 | - | 79 | 79 | 79 | 78 | 79 |
| GmNADP-ME2.1 | 81 | 82 | 79 | - | 98 | 82 | 81 | 81 |
| GmNADP-ME2.2 | 81 | 81 | 79 | 98 | - | 81 | 81 | 81 |
| GmNADP-ME3.1 | 77 | 77 | 79 | 82 | 81 | - | 92 | 89 |
| GmNADP-ME3.2 | 80 | 80 | 78 | 81 | 81 | 92 | - | 98 |
| GmNADP-ME3.3 | 77 | 77 | 79 | 81 | 81 | 89 | 98 | - |

**Table S1 Comparison of amino acid sequences of Arabidopsis, castor, and soybean MEs.** Sequences were obtained from Phytozome (<https://phytozome-next.jgi.doe.gov/>) and multiple alignments were performed with Clustal W (<https://www.genome.jp/tools-bin/clustalw>). Values corresponded to percentages of identity: >70% highlighted in dark grey, 60-70% highlighted in light grey, and <42% unhighlighted. ^a^characterized in this work.

| **Entry** | **Model** | **Root mean squared deviation** | **Template modeling score** | **Identity (%)** | **Sequence length** | **Equivalent residues** |
| --- | --- | --- | --- | --- | --- | --- |
| **NAD-ME2.4** | AF-I1KIF0-F1-model_v4.pdb | - | - | - | 604 | - |
| **NAD-ME2.3** | AF-I1JKC3-F1-model_v4.pdb | 0.36 | 0.97 | 97 | 604 | 585 |
| **NAD-ME2.1** | AF-I1L6N5-F1-model_v4.pdb | 0.72 | 0.96 | 86 | 601 | 585 |
| **NAD-ME1** | AF-I1JMI9-F1-model_v4.pdb | 2.1 | 0.94 | 65 | 622 | 604 |

**Table S2 Structural aligment of soybeanNAD-ME models.** Models were obtained from AlphaFold Protein Structure Database (<https://alphafold.ebi.ac.uk/>) and aligned withNAD-ME2.4 using Pairwise Structural Aligment Tool (<https://www.rcsb.org/alignment>).

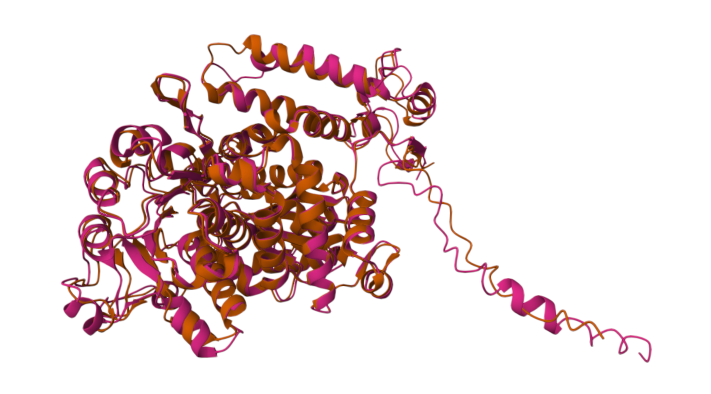

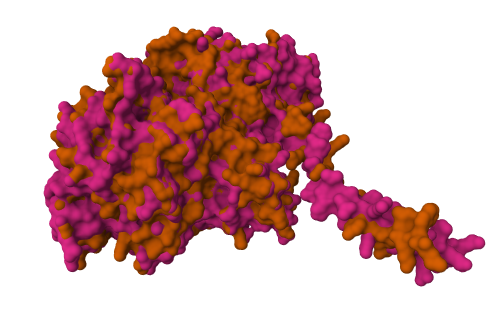

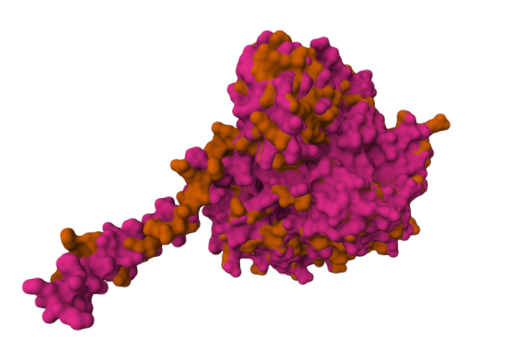

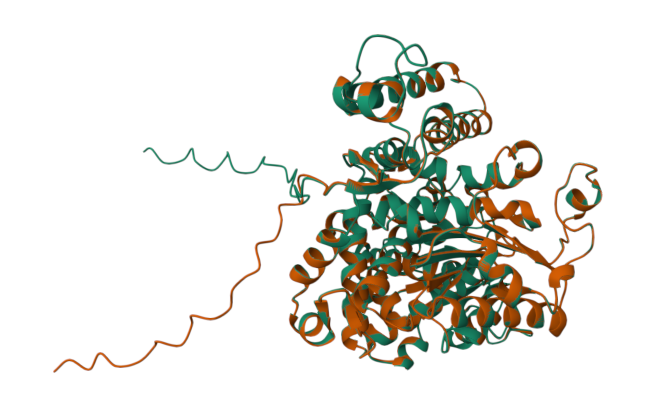

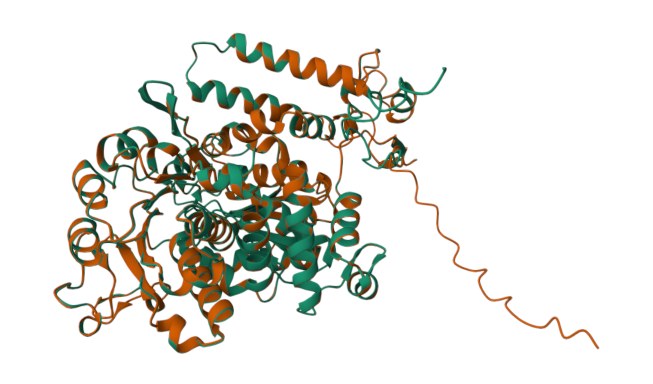

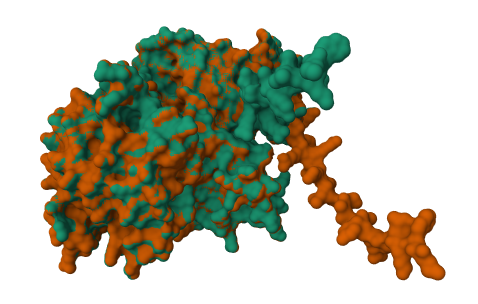

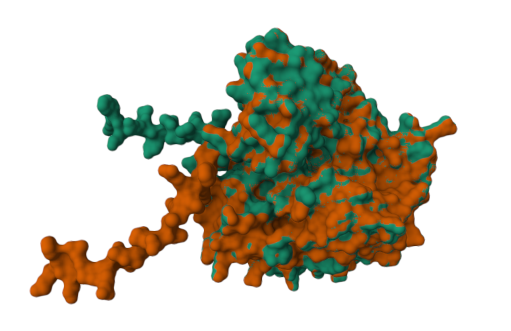
**A**

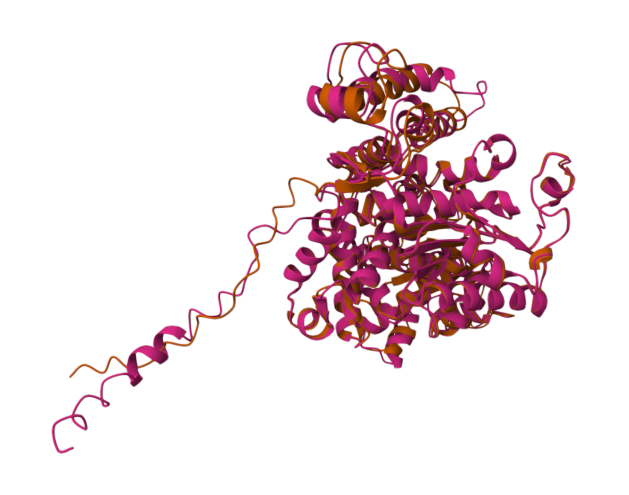
**B**

**Figure S1 Alignment of structural models of soybean NAD-ME2.4 with NAD-ME2.3 (A) or NAD-ME1 (B).** Two views (vertical rotation of 180°) are shown for each surface and ribbon representation. NAD-ME2.4 is indicated in orange, while NAD-ME2.3 and NAD-ME1 are shown in green and pink, respectively.

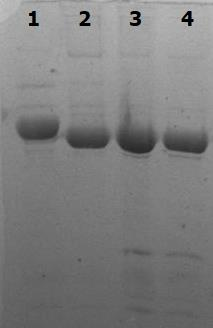

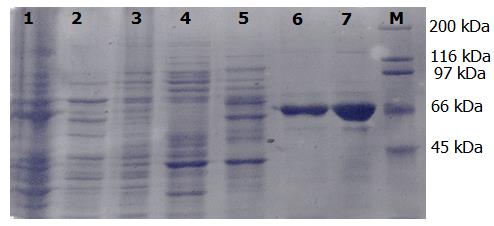

**97 kDa**

**66 kDa**

**A**

**B**

**Figure S2 A)** SDS-PAGE of the purification steps of recombinant NAD-ME1. M: molecular mass markers; lanes 1 and 2: insoluble and soluble fractions of the bacterial extract, respectively; lane 3: fraction not retained on the Ni-NTA column; lanes 4 and 5: fractions collected after the passage of binding and washing buffers; lane 6: eluted fraction; lane 7: concentrated protein. **B)**Similar results were obtained for the purification of the other isoforms: NADP-ME1.1 (lane 1), NAD-ME2.1 (lane 2), NAD-ME2.3 (lane 3), and NAD-ME2.4 (lane 4). In each lane, 4 µg of each concentrated and purified protein was loaded.

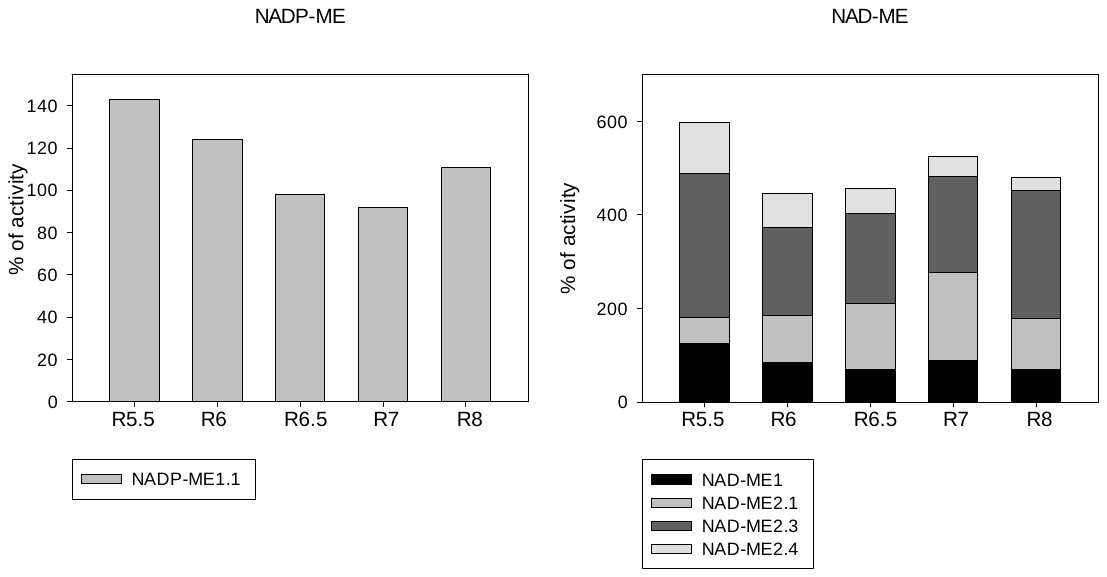

*

*

*

*

*

*

*

*

*

*

*

*

**A	B**

**Figure S3 Activity of the NADP- (A) and NAD-ME (B) isoforms in the presence of mixtures mimicking cellular conditions of embryos at different developmental stages.** Enzymatic activity was evaluated in the presence of total polar metabolite extracts from soybean embryos on R5.5-R8 stages. Data are presented as ratio (in percentages) of activity compared to a control without extracts (100%; the asterisk indicates a significant difference in Student's t-test, *p* < 0.05). Values are the mean of triplicate determinations with standard deviation less than 20%.

See file

**Video S1 3D alignment of NAD-ME2.4 (in orange) with NAD-ME2.3 (in green).**

See file

**Video S2 3D alignment of NAD-ME2.4 (in orange) with NAD-ME1 (in pink).**
